# HBMITool: a user-friendly software for labeling Human Brain Microscopy Images

**DOI:** 10.1101/699215

**Authors:** Hungju Wang, Lea T Grinberg, Maryana Alegro

## Abstract

One of the most popular tools for quantifying protein expression is Immunofluorescence (IF). Although IF is widely applied in drug discovery research and assessing disease mechanisms, it has great room for improvement on the task of analyzing human postmortem brain samples. IF analysis of postmortem human tissue relies mostly on manual interaction, which is often error-prone and leading to low inter and intra-observer reproducibility. The high level of autofluorescence caused by accumulation of lipofuscin pigment during aging impedes systematic analyses of human postmortem brain samples. A method for automating cell counting and classification in IF microscopy of human postmortem brains was proposed before, which speeds up the quantification task while improving reproducibility. To correct for misclassified cells by the algorithm, we created HBFMTool, a software package that ease the process of editing the result produced by cell detection/classification algorithm.

## I. INTRODUCTION

Target validation and cell selection provide critical insights into mechanisms driving disease, particularly at the early steps in the drug discovery process. Immunofluorescence (IF) microscopy applies antibodies and other affinity reagents with fluorescent proteins on tissue and may unlock information into molecular cascades while preserving the tissue integrity. IF is particularly important in quantifying protein expression and understanding cell function. IF experiments often occur in big batches resulting in high amounts of data, making manual analysis very time-consuming and prone to error including high inter and intra-observer variability and low reproducibility [1].

A computer-based segmentation method was proposed that is suitable for handling IF microscopy from our thick postmortem human brain sections, aiming to improve user productivity by reducing the counting time and also reducing inter and intra-observer variability [2]. Our proposal consists in applying dictionary learning and sparse coding for cell modeling. Dictionary learning is a cutting-edge method for learning signal representations that better fit underlying data, thus improving description [3], while sparsity allows better portrayal of salient image properties and produces features more likely to be separable, which is useful for machine learning [3, 4].

In order to correct for misclassified cells and improve the cell detection algorithm, HBFMTool, a user-friendly, general-purpose, editing/labeling software is proposed. The overall work flow of the software is as follows: (1) run the HBFMTool MATLAB script that performs automated cell detection on all the raw IF sample images, (2) run the HBFMTool graphical user interface and start the correcting process, (3) save the edited result via the graphical user interface. The software records and saves the algorithm-labeled cell locations, the human-labeled cell locations, the different categories of cells and the number cells in each category. By having all these data, HBFMTool facilitates further training of the automated cell-detection algorithm and result collection and analysis for neuroscientists.

## II. MATERIALS AND METHOD

**Fig. 1.**
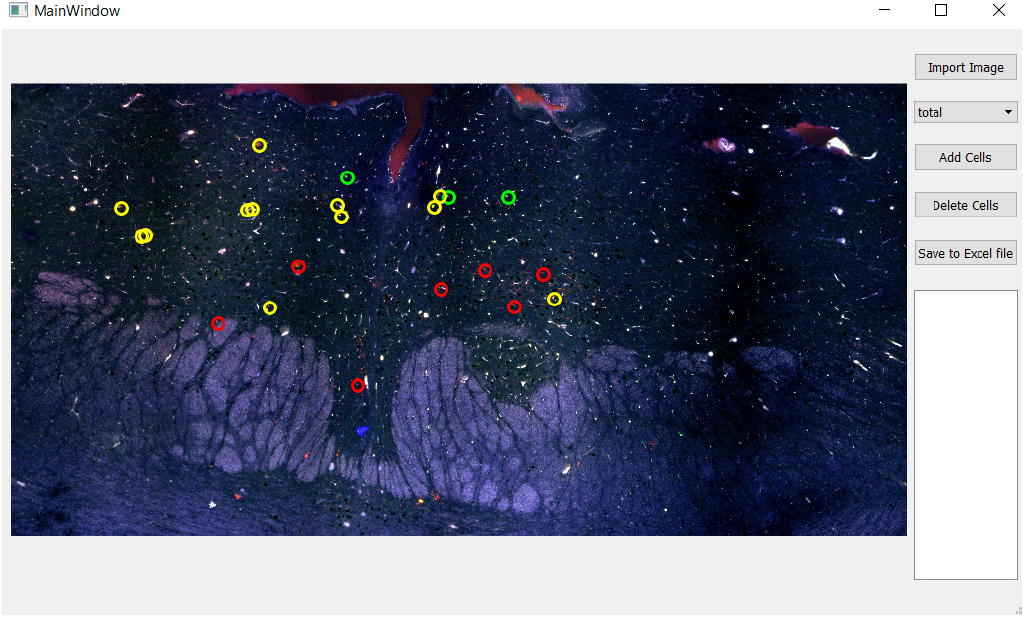
HBFMTool: graphical user interface with customized marks

This software is developed on a Windows 10 platform but works on both Windows and Linux Ubuntu. Since the output of the cell detection algorithm is binary images of different kinds of cells, we first take theses binary images and use the region prop function from MATLAB to compute the coordinates of auto-detected cells with respect to the coordinate frame of each binary image. We then save it in an excel file along with other important information, such as the number of cells in each binary image. One thing to note is that the dimension of the original IF image is four times larger than the binary image. Thus, in the next step when we overlay customized marks, which represents the cells, with the original IF image, we have to multiply the coordinates of the recorded cells by four.

Once the pre-detected cells are overlaid, the user can see the original image with marks and thus, able to visually correct for mislabeled cells. In the event of cells that are not detected by the algorithm, the user can first select the category and overlay different colors of marks on the image. On the other hand, when the algorithm misclassified a cell, the user can click the delete button and the mark will disappear. Currently, the way HBFMTool keeps track of the coordinates is by having three different lists, each of them corresponds to a category of cells.

## III. Results/Discussion

With the above implementation of the software, HBFM-Tool comes out to be an user-friendly, general purpose and easily-adjustable tool. It allows neuroscientists to examine and correct the result of the cell detection algorithm. It also allows the algorithm to take into the feedback of the neuroscientists and potentially improve over time.

